# Temporal structure of chemical stress controls single-cell inhibition and recovery in photosynthetic microorganisms

**DOI:** 10.64898/2025.12.22.695903

**Authors:** Linhong Xiao, Adeolu Ogunleye, Joëlle Rüegg, Yuan Cui, Lars Behrendt

## Abstract

Although fluctuating conditions are a hallmark of microbial habitats, how the timing of chemical stress influences single-cell inhibition and recovery is not well resolved. Using a microfluidic platform, we continuously tracked photosynthetic performance in thousands of individual cells of a coral-associated microalga (*Symbiodiniaceae* sp.) exposed to identical cumulative doses of the photosystem II inhibitor diuron delivered either as constant or fluctuating profiles. Fluctuating exposure produced stronger early inhibition than constant exposure, but as concentrations declined it enabled partial recovery that did not occur under time-averaged constant conditions. These dynamics revealed distinct response subpopulations that differed in both the magnitude and timing of inhibition, including groups that regained activity exclusively under fluctuating stress. Quantifying per-cell decline kinetics showed that fluctuating exposure synchronizes the onset of inhibition across cells, creating a narrow temporal window in which a subset of cells can recover once stress levels fall, whereas constant exposure yields more heterogeneous but uniformly declining trajectories. These results demonstrate that stress timing, not cumulative dose alone, governs whether photosynthetic inhibition is reversible at the single-cell level. More broadly, our findings illustrate how temporal variability and intrinsic phenotypic heterogeneity jointly govern cellular function, highlighting stress timing as an important and often overlooked axis shaping microbial performance in dynamic environments.

**Significance statement:** Environmental stress in nature is rarely constant; instead, it fluctuates over minutes to days. Yet most laboratory assays rely on static exposures, leaving the role of stress timing poorly understood. Using microfluidics, we imposed precisely timed chemical stress on individual cells of a coral-associated microalga and monitored their photosynthetic performance continuously. Fluctuating exposure caused strong initial inhibition but later allowed partial recovery in a subset of cells, revealing functional heterogeneity that constant stress concealed. These results show that when stress occurs, its temporal structure can be as important as how much stress is delivered. By uncovering mechanisms that govern reversibility and heterogeneity in a key coral symbiont, this work provides a mechanistic basis for understanding microbial responses in dynamic environments.

## Introduction

Microorganisms rarely encounter environmental stress in a constant form. Natural fluctuations in temperature, chemistry, and resource availability impose temporal patterns that can strongly influence how individual cells respond. Yet the consequences of this temporal structure for microbial dynamics remain poorly resolved, even though stress timing and intrinsic cellular heterogeneity could shape population outcomes in ways that cumulative dose alone does not predict.

Although environmental fluctuations are widespread, how their timing influences cellular responses and how these responses scale to affect higher-level processes remain open questions. These issues are particularly relevant for photosynthetic microorganisms, where individual cell-level photophysiology strongly shapes population performance and contributes to broader ecological function. Coral reef symbioses exemplify this connection: corals rely on dinoflagellate symbionts of the family Symbiodiniaceae, whose single-cell photophysiology determines host productivity and reef resilience (1, 2). This association is highly sensitive to environmental perturbations that can trigger coral bleaching (3, 4), the breakdown of the partnership between host and symbiont. However, it remains unclear how temporal variability in environmental stress, such as fluctuating chemical exposure, shapes the balance between inhibition and recovery in Symbiodiniaceae cells. Clarifying this relationship is essential for understanding how temporal stress structure affects the cellular processes on which higher-level ecological responses depend.

Controlled chemical perturbations provide a powerful way to dissect how temporal variability shapes cellular stress responses. To investigate this question in an ecologically grounded context, we focused on diuron (3-(3,4-dichlorophenyl)-1,1-dimethylurea; DCMU), a well-characterized photosystem II inhibitor (5, 6) and a coral-relevant herbicide (7–9). Field surveys show that diuron concentrations in reef waters fluctuate markedly over time (10, 11), likely exposing symbiotic algae to dynamic rather than static chemical regimes. Such fluctuations create an opportunity to test how the timing of chemical stress -independent of cumulative dose- shapes inhibition and recovery trajectories in individual *Symbiodiniaceae* cells, and how temporal variation generates the functional heterogeneity that underlies divergent cell outcomes.

We hypothesized that the temporal exposure structure interacts with single-cell heterogeneity to produce distinct response subpopulations with different inhibition and recovery trajectories. Addressing this hypothesis requires precisely timed perturbations and continuous single-cell measurements of photosynthetic performance. Recent advances in microfluidic control (12–14) and single-cell photophysiology (15–17) now make this possible, enabling dynamic chemical profiles to be applied while the same cells are monitored in near real time. Here, we used this framework to expose thousands of Symbiodiniaceae cells to either constant or fluctuating diuron profiles that were matched for cumulative dose. We show that temporal structure, not total dose, governs the onset timing and reversibility of photosynthetic inhibition, revealing distinct subpopulations that emerge specifically under fluctuating stress. Together, these findings establish how temporal variability interacts with intrinsic phenotypic heterogeneity at the single-cell level, providing a mechanistic foundation for understanding how environmental fluctuations shape microbial stress responses.

### Temporal variability in chemical stress shapes inhibition and recovery dynamics

Individual Symbiodiniaceae cells were confined in an array of microwells integrated into a microfluidic chip (Figure 1a–b), which enabled controlled delivery of chemical stress and continuous microscopic monitoring of hundreds of individual cells over long experimental timescales. The cells were exposed either to steady diuron concentrations or to fluctuating diuron concentrations following a 24 h negative-cosine waveform (Figure 1c, S1). Fluorescein tracing confirmed that the realized waveform closely matched the imposed profile (R² = 0.996; Figure S2). Each individual cell was measured, in darkness, every 15 minutes over a 24 h period using a pulse-amplitude-modulated (PAM) chlorophyll fluorometry microscope. This generated single-cell trajectories of the dark-adapted maximum quantum yield of photosystem II (*F_v_*/*F_m_*), a widely accepted indicator of photosynthetic performance.

**Figure 1.**
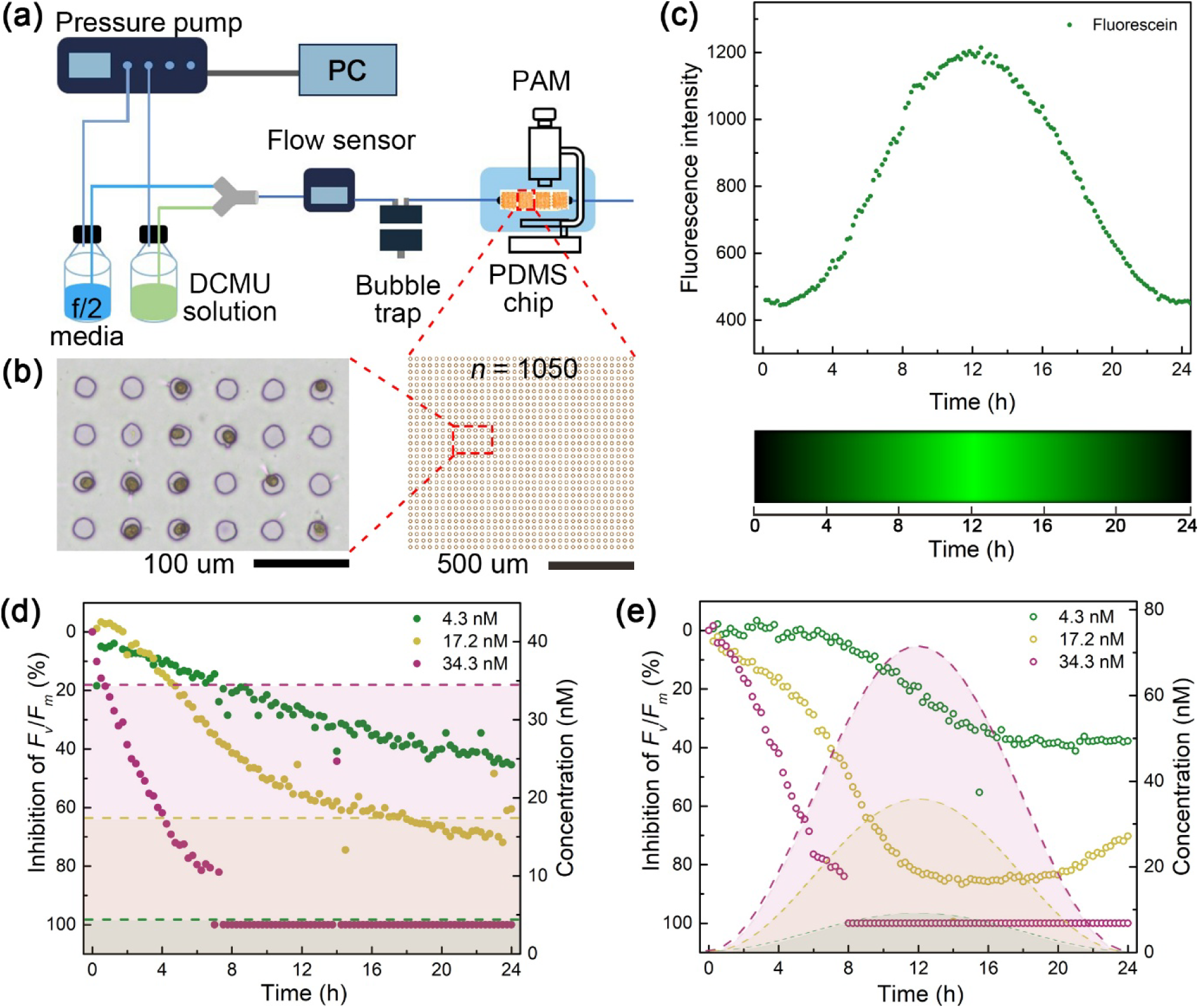
Microfluidic platform for applying steady and fluctuating diuron exposures to individual Symbiodiniaceae cells. **(a)** Schematic of the microfluidic setup used to deliver user-defined diuron profiles. A pressure-driven pump system mixed f/2 medium with diuron solution at controlled ratios, and flow was monitored by inline sensors before entering a bubble trap and the PDMS chip positioned under a PAM chlorophyll fluorometry microscope. This configuration enabled continuous, single-cell measurements of *F_v_*/*F_m_* under either steady or fluctuating exposure regimes. **(b)** Left: Bright-field image of microwells (20 µm diameter) used to immobilize individual cells. Right: Layout of the microwell array (*n* ≈ 1050 per field of view). **(c)** Fluorescein measurements confirming that the realized fluctuating profile closely matched the programmed negative-cosine waveform with a peak at 12 h. Bottom: kymograph of fluorescein intensity over 24 h. **(d)** Inhibition of *F_v_*/*F_m_* over time for cells exposed to steady diuron concentrations at 4.3, 17.2, and 34.3 nM. Points show mean inhibition across individual cells (*n* = 539, 231, and 557). Shaded regions denote the corresponding steady concentration levels (right axis). (e) Inhibition of *F_v_*/*F_m_* under fluctuating diuron exposure at the same time-averaged concentrations (*n* = 411, 371, and 518). Shaded regions show the fluctuating concentration profiles. Steady and fluctuating treatments delivered the same cumulative dose over 24 h.

To isolate the effect of stress timing from total chemical dose, the time-averaged concentration in each fluctuating treatment was matched to the corresponding steady treatment, meaning both delivered the same overall amount of diuron (the area under the concentration–time curve) over a 24 h period. The three concentrations used were 4.3 nM, 17.2 nM, and 34.3 nM. These concentrations were selected to span a range from mild to strong photophysiological inhibition while overlapping with the lower to intermediate range of diuron concentrations reported from nearshore coral reef environments, where exposure typically occurs as transient pulses rather than constant plateaus. Because fluctuating profiles involve programmed rises and falls in concentration generated by dynamically varying the ratio of diuron-containing and diuron-free media, peak concentrations for the fluctuating treatments were 9.2 nM, 35.8 nM, and 71.5 nM, respectively. All reported reductions in *F_v_*/*F_m_* are relative to unexposed controls and were conducted across a minimum of two independent experiments. Under steady exposure, inhibition of *F_v_*/*F_m_* increased with both concentration and duration (Figure 1d; Table S1): 4.3 nM reduced mean *F_v_*/*F_m_* by 26.3 ± 18.3% (± standard deviation) after 12 h and 45.3 ± 19.4% after 24 h (*n* = 539); 17.2 nM reduced the mean by 55.7 ± 30.8% after 12 h and 60.5 ± 3.1% after 24 h (*n* = 231); 34.3 nM fully inhibited activity within 7 h (*n* = 557).

At equivalent time-averaged concentrations, fluctuating exposures caused sharper but partly reversible inhibition (Figure 1e; Table S1). At the lowest exposure of 4.3 nM (peak: 9.2 nM), diuron caused photoinhibition of 19.2 ± 3.6% after 12 h and 37.7 ± 27.4% after 24 h (n = 511). At 17.2 nM (peak: 35.8 nM), inhibition rose rapidly to 82.3 ± 11.0% after 12 h and then partially relaxed to 70.2 ± 11.8% by 24 h (*n* = 371; Figure S3), indicating partial recovery within the experimental window. The highest diuron concentration, 34.3 nM (peak: 71.5 nM), caused complete inhibition within 8 h (*n* = 518), with no recovery observed over the duration of the experiment. Together, these observations indicate that fluctuating exposure alters how inhibition accumulates over time. At intermediate concentrations, fluctuating stress can initially amplify inhibition relative to steady exposure delivering the same overall amount of diuron, yet later permit partial recovery as concentrations decline (Figure 1e, Figure S3). Thus, temporal structure does not simply modulate the severity of inhibition, but reshapes the relationship between delivered exposure and cellular response over time.

### Timing influences inhibition dynamics when total dose is the same

To further assess how exposure timing influences inhibition dynamics, we analyzed the cumulative diuron dose experienced by cells under both steady and fluctuating conditions (Figure 2a).

**Figure 2.**
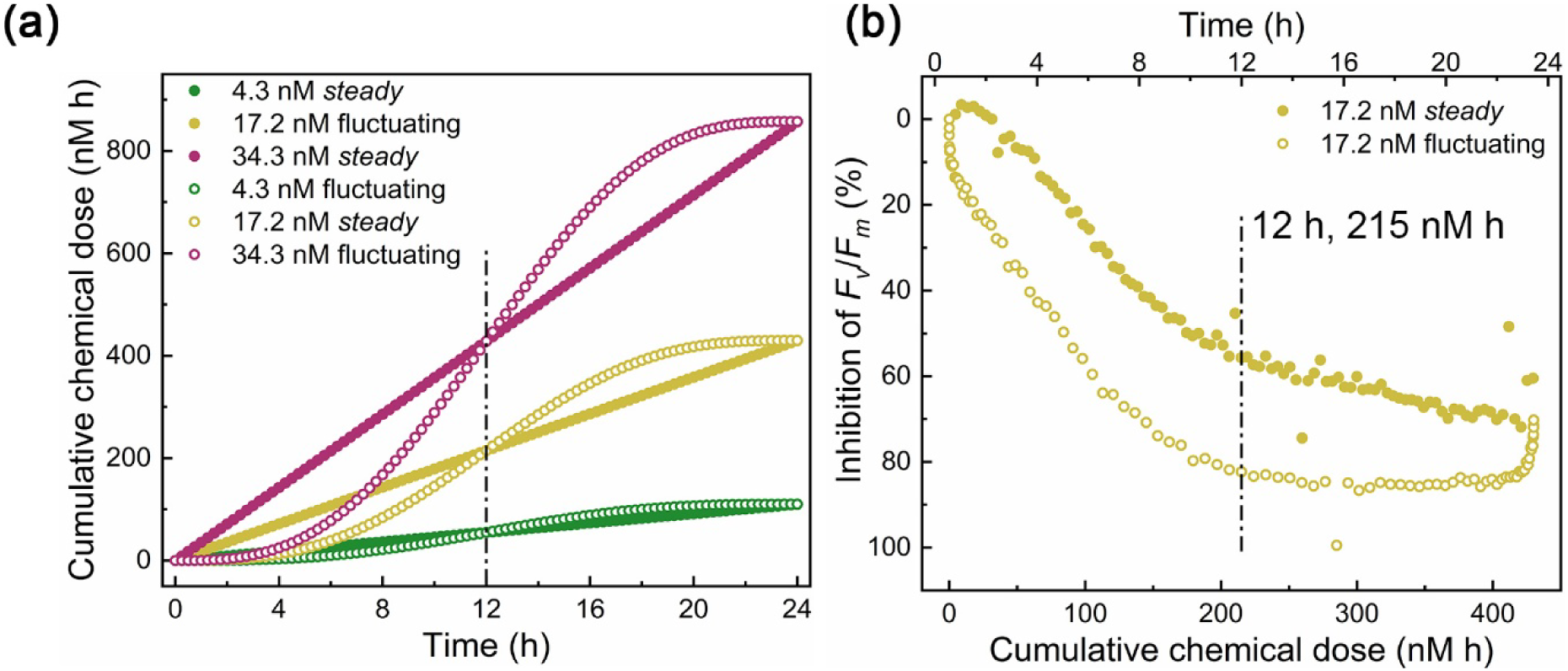
Photophysiological responses as a function of cumulative diuron dose. **(a)** Cumulative diuron dose (area under the concentration–time curve) for steady (solid symbols) and fluctuating (open symbols) exposure regimes at 4.3, 17.2, and 34.3 nM (colors as shown). Minor deviations (2–4%, Figure S3) between profiles reflect small differences in realized mixing ratios. **(b)** Inhibition of *F_v_*/*F_m_* as a function of cumulative dose for the 17.2 nM treatment. Steady (solid) and fluctuating (open) trajectories diverge at identical cumulative doses, indicating that timing, not total dose, drives differences in photo-physiological response. The dashed vertical line marks the shared cumulative dose at 12 h (≈215 nM·h). Corresponding analyses for 4.3 and 34.3 nM exposures are shown in Figure S3.

Cumulative dose *D* was defined as the area under the concentration–time curve, D = ∫ₜ₀^ₜₓ c(t) · dt, where *c(t)* is the diuron concentration at time *t*, and *t₀* and *tₓ* mark the start and end of the exposure window, yielding an integrated exposure in nM·h. Although both regimes are designed to deliver identical cumulative doses at 12 h and 24 h, minor deviations of 2–4% (Figure 2a, S3a) were observed, arising from small differences in microfluidic mixing ratios. These deviations were insufficient to account for the pronounced differences in inhibition dynamics. Thus, differences in inhibition reflect the timing and structure of exposure rather than differences in total delivered dose. At intermediate diuron concentrations (17.2 nM average; peak 35.8 nM), both regimes delivered approximately 215 nM·h by the 12 h midpoint (Figure 2b). Despite this equivalence, cells under fluctuating exposure showed significantly stronger inhibition at 12 h than those under steady conditions (82.3 ± 11.0% vs. 55.7 ± 30.8%; Mann-Whitney U test, *p* < 10⁻⁴, Figure 2b, S3c), consistent with early peak concentrations in the fluctuating profile. During the second half of the experiment, inhibition under the fluctuating regime partially relaxed as concentrations declined, whereas steady exposure produced a slower, continuous decrease. By 24 h, both treatments converged to nearly equivalent inhibition levels (70.2 ± 11.8% vs. 60.5 ± 3.1%; Mann-Whitney U test, *p* < 10⁻⁴), highlighting that fluctuating exposure primarily alters the timing -not the final magnitude- of inhibition. At lower and higher concentrations, timing effects diminished. At 4.3 nM (peak: 9.2 nM), inhibition at 24 h did not differ significantly between fluctuating and steady exposures (37.7 ± 27.4% vs. 45.3 ± 19.4%; Mann-Whitney U test, *p* = 0.82; *n* = 411 vs. 539; Figure S3b). At 34.3 nM (peak: 71.5 nM), both regimes caused complete inhibition within 8 h (*n* = 518 and 557; Figure S3d), indicating that under extreme stress intensities, exposure pattern no longer alters the outcome. Thus, timing effects emerged most clearly at intermediate stress, where partial recovery was observed within the experimental window. Together, these results show that under moderate stress, temporal structure strongly shapes photosynthetic function beyond what is predicted from cumulative dose alone.

### Fluctuating stress reveals distinct functional subpopulations

To examine how exposure timing shaped population structure, single-cell *F_v_*/*F_m_* trajectories were grouped by K-means clustering. Distinct response clusters were identified under both steady and fluctuating regimes across diuron concentrations (Figure 3a–d; Figures S4–S5). At 17.2 nM, clustering revealed pronounced functional diversification, including subpopulations that maintain activity, lose function rapidly, or partially recover after peak stress under fluctuating exposure. For example, under steady diuron exposure, cells in cluster 2 maintained an average *F_v_*/*F_m_* of 0.48 ± 0.05 (*n* = 40; representing 17% of the population) after 12 h, whereas those in cluster 7 (*n* = 42; 18% of the population) were fully inhibited (Figure 3a and c). Under fluctuating conditions, additional subpopulations emerged that exhibited partial recovery as concentrations declined. One such group (cluster 2; *n* = 34; 9% of the population) maintained *F_v_*/*F_m_* = 0.15 ± 0.05 at the 12 h peak, while another (cluster 7; *n* = 58; 18% of the population) declined to <0.05 ± 0.02 before regaining activity during the final 4 h of exposure (Figure 3b and d). Comparable patterns were observed at lower and higher diuron concentrations (Figures S4–S5). For instance, under fluctuating conditions at 4.3 nM, cells in cluster 2 (*n* = 53; 12% of the population) showed minor inhibition, whereas those in cluster 5 (*n* = 32; 7% of the population) were more sensitive. Together, these results show that fluctuating exposure reshapes how cells are distributed across functional trajectories, making recovery, rare or absent under steady exposure, more common when stress levels decline. This supports the hypothesis that temporal structure interacts with intrinsic heterogeneity to generate distinct single-cell outcomes.

**Figure 3.**
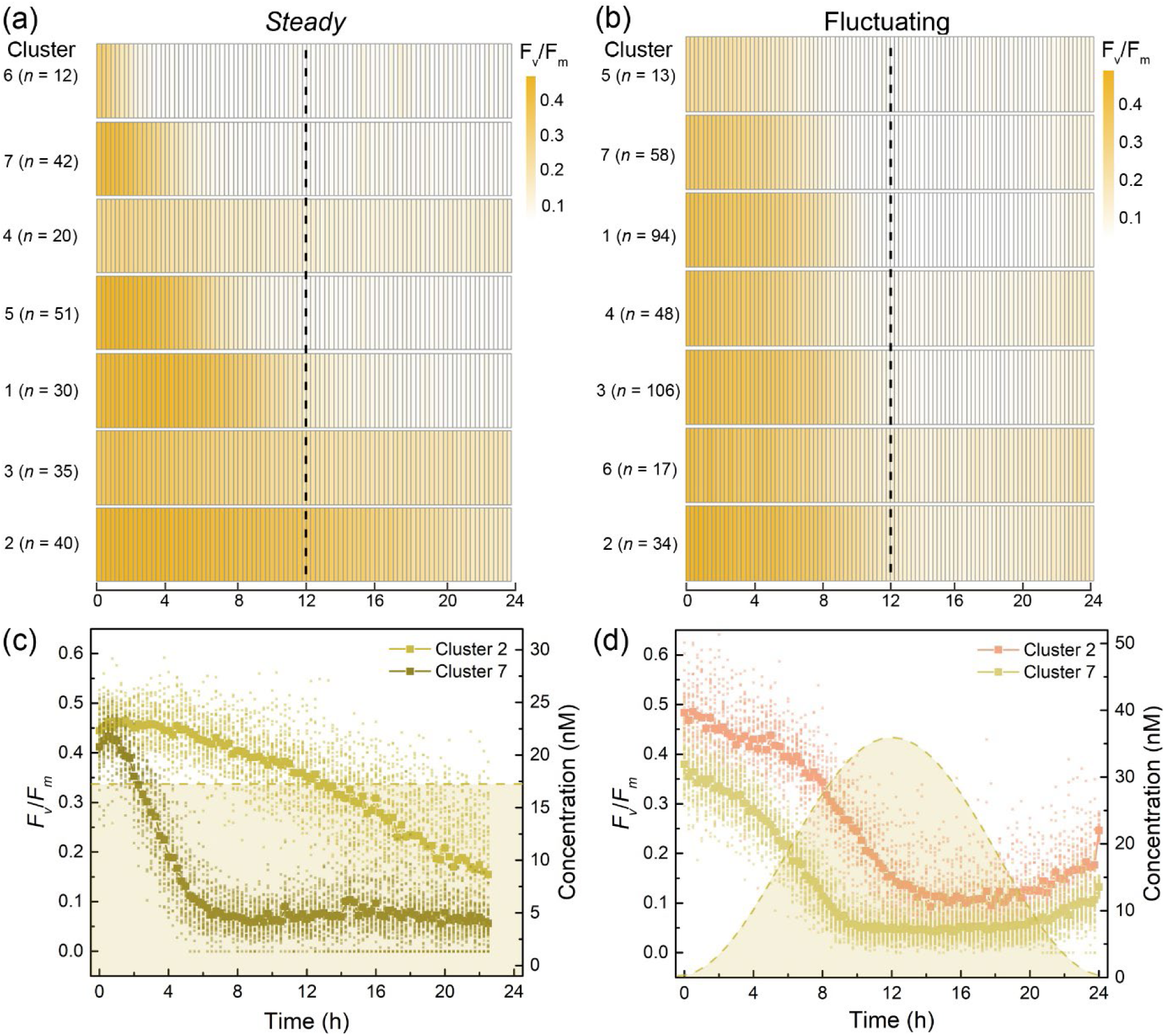
Distinct subpopulation dynamics under steady and fluctuating diuron exposures. **(a, b)** Heatmaps of cluster-averaged *F*ᵥ/*F*ₘ trajectories for *Symbiodiniaceae* subpopulations exposed to steady (a) or fluctuating (b) diuron regimes at 17.2 nM (time-averaged). Clusters were identified by K-means using individual cell trajectories (see Methods). Numbers indicate cells per cluster. **(c, d)** Representative trajectories from two clusters illustrating contrasting functional responses. Cluster 2 cells maintained higher *F*ᵥ/*F*ₘ, whereas cluster 7 cells showed stronger inhibition. Under fluctuating exposure (d), some cells in clusters 2 and 7 regained activity as concentrations declined, whereas no comparable recovery occurred under steady exposure (c). These patterns show that fluctuating stress reveals recovery-associated trajectories that remain absent or rare under constant exposure.

### Cells vary widely in how fast they lose photosynthetic activity

To quantify how rapidly individual cells lost photosynthetic function, we calculated a half-life (t₁/₂) for each single-cell trajectory, defined as the time at which *F*ᵥ/*F*ₘ declined to half of its initial value (Figure 4a). Across all treatments, t₁/₂ values exhibited broad cell-to-cell variability, consistent with heterogeneous stress responses. At the lowest diuron concentration (4.3 nM), t₁/₂ distributions differed modestly between exposure regimes, with fluctuating exposure yielding slightly longer median t₁/₂ values than steady exposure (median 11.64 h vs. 10.85 h; Mann–Whitney U test, *p* = 0.0017; *n* = 410 and 538, respectively; Figure 4b). This indicates that when inhibition was mild, exposure timing had only a modest effect on decline kinetics. At the intermediate concentration (17.2 nM), median t₁/₂ values were nearly identical under steady and fluctuating exposure (6.87 h vs. 6.95 h; Mann–Whitney U test, *p* = 0.91; *n* = 230 and 370). However, the spread of t₁/₂ values differed markedly between regimes. Under fluctuating exposure, t₁/₂ values were tightly distributed (interquartile range 1.90 h), whereas steady exposure produced a much broader distribution with a pronounced long tail (interquartile range 5.45 h; Figure 4b). Thus, although the typical time to half-inhibition was similar, fluctuating exposure synchronized the onset timing of functional decline across cells, while steady exposure allowed cells to cross the half-activity threshold according to individual sensitivities. At the highest concentration (34.3 nM), fluctuating exposure again produced significantly longer t₁/₂ values than steady exposure (median 3.71 h vs. 2.74 h; Mann–Whitney U test, *p* < 0.0001; *n* = 517 and 556), indicating that exposure timing strongly modulated decline kinetics under severe stress. Although t₁/₂ does not capture recovery directly, the synchronized decline observed under fluctuating exposure helps explain why recovery-associated trajectories emerged predominantly in this regime. This is because cells reached strong inhibition within a narrow temporal window and a subset was positioned to regain activity once stress levels declined. Under steady exposure, by contrast, cells entered inhibition over a broad range of times and continued to decline, suppressing recovery. Together, these results show that stress timing shapes not only the extent of inhibition but also when cells lose function, thereby influencing which subpopulations can recover as conditions improve.

**Figure 4.**
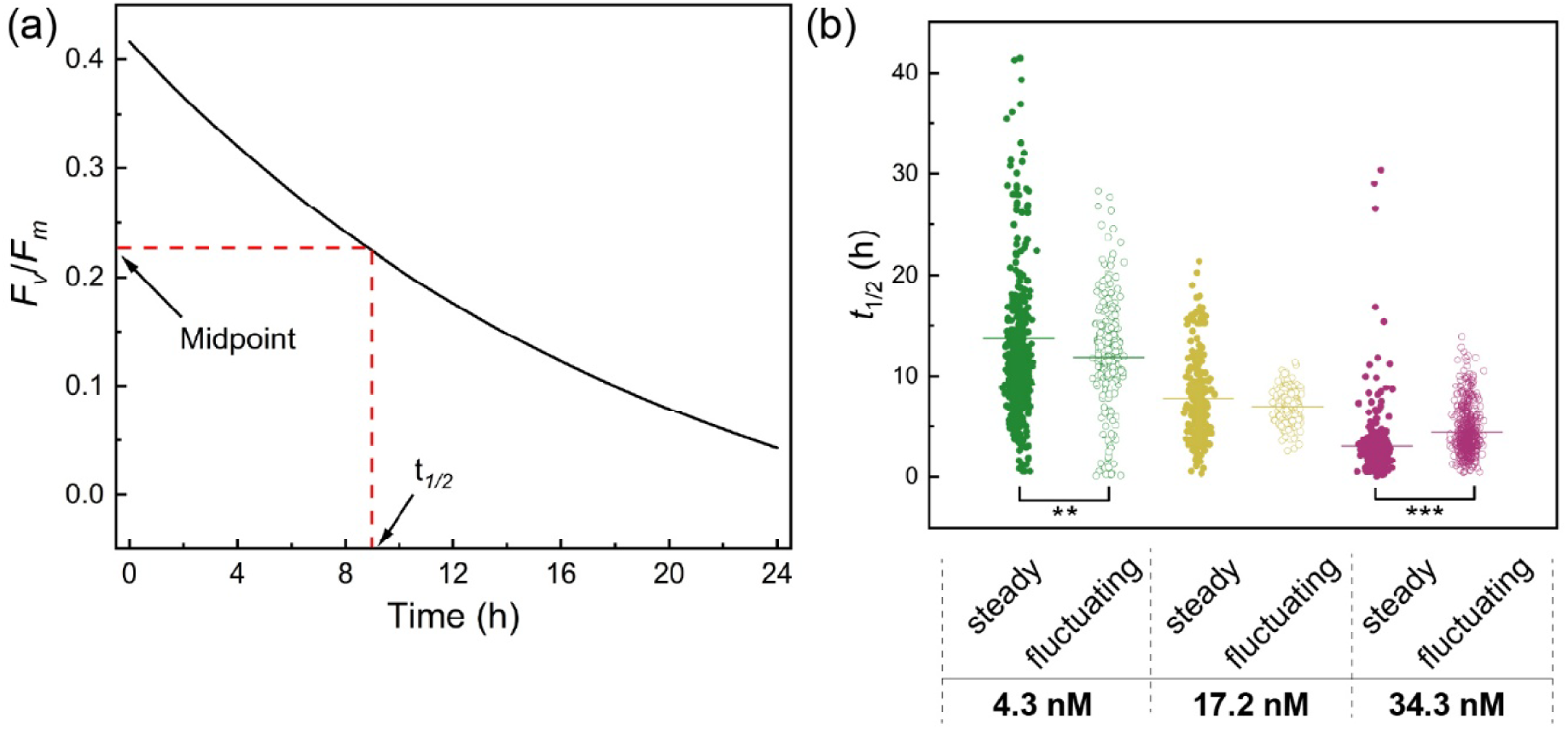
Single-cell decline kinetics (t₁/₂) under steady and fluctuating diuron exposure. **(a)** Illustration of the half-life metric (t₁/₂), defined as the time when a cell’s *F*ᵥ/*F*ₘ first falls to 50% of its initial value. **(b)** Distribution of t₁/₂ values across cells exposed to steady (solid symbols) and fluctuating (open symbols) diuron at 4.3, 17.2, and 34.3 nM. Horizontal bars denote median values. At intermediate stress levels (17.2 nM), median t₁/₂ values were similar between exposure regimes, but fluctuating exposure yielded a markedly narrower distribution, indicating synchronized onset of inhibition, whereas steady exposure produced a broader spread of decline kinetics. At low and high diuron concentrations, median t₁/₂ values differed significantly between regimes. Asterisks indicate statistical significance of pairwise comparisons (Mann–Whitney U test; **p < 0.01; ***p < 0.001).

## Discussion

### Temporal structure of stress as a determinant of microbial resilience

Our findings show that how stress is delivered, not simply how much, is a key determinant of photophysiological outcomes in individual Symbiodiniaceae cells. Fluctuating diuron exposures produced strong transient inhibition of PSII but, at intermediate concentrations, were associated with partial recovery within the experimental window, whereas time-averaged constant exposures did not show comparable recovery over the same period. These observations indicate that the waveform and timing of stress exposure redirect the trajectory of cell inhibition and recovery over time, revealing outcomes that are not predictable from cumulative dose alone.

Environmental fluctuations are ubiquitous: temperature, chemical inputs, and nutrient availability vary across diel and episodic timescales in most ecosystems. Prior work has shown that such variability can profoundly alter microbial growth, stress tolerance, and adaptation (6, 13, 18). Likewise, short pulses of herbicides in natural waters can elicit physiological effects distinct from constant exposures despite identical integrated doses (19–21). However, these studies have generally applied static or step-change herbicide profiles, leaving the specific role of exposure waveform unresolved. Recent work demonstrates that fluctuating environments can give rise to physiological states that are not captured by steady-state responses, because transient responses shape downstream behavior (13). Our findings parallel this: the early inhibitory peak in the fluctuating profile subjected all cells to severe stress simultaneously, synchronizing their initial decline, which in turn alters subsequent cell trajectories and produces inhibition–recovery dynamics that were not observed under constant exposure within the same timeframe. By disentangling temporal structure from cumulative dose, our study adds a mechanistic dimension that standard static assays do not capture.

#### Phenotypic heterogeneity buffers populations against fluctuating stress

Only a subset of cells recovered following peak stress, indicating that intrinsic single-cell heterogeneity is a major driver of population-level resilience. Clustering revealed subpopulations that either maintained activity, lost function rapidly, or regained PSII performance once stress declined. Calculated single-cell half-life times quantified these differences, demonstrating that cells vary widely in how long they retain function under identical cumulative doses. This heterogeneity resembles persister-like dynamics in bacterial populations exposed to antibiotics and other inhibitors (22, 23), and parallels observations from fluctuating nutrient and environmental regimes (12, 13). Our observation that recovery-associated trajectories emerge preferentially under fluctuating exposure profiles aligns with the idea that temporal variability amplifies the functional consequences of intrinsic phenotypic differences, an effect also seen in systems where fluctuation-adapted states emerge under rapid environmental switching (13). It is important to acknowledge that we did not track individual cell-cycle or life-stage identities in Symbiodiniaceae; such physiological states almost certainly influence how cells withstand and recover from stress and may underlie part of the observed heterogeneity.

#### A mechanistic link between environmental variability and resilience

Establishing the role of both temporal structure and phenotypic heterogeneity carries broader implications for microbial ecology and coral symbiosis. In reefs, symbiont photosynthetic performance is tightly coupled to host health, and the persistence of even a minority of functional cells may delay or modulate bleaching responses. Previous work shows that low-level herbicides can induce reversible inhibition (19) and that chronic diuron exposure interacts with elevated temperature to lower bleaching thresholds and reduce photosynthetic performance, calcification, and survival in corals and other symbiotic taxa (5, 10, 20). Our results extend this work by showing mechanistically that temporal exposure structure alone, independent of cumulative dose, can shape whether functional subpopulations persist over time, even when overall inhibition levels converge.

Because the fluctuating regime imposes a sharp peak early in the exposure window, all cells encounter severe inhibition at nearly the same moment, synchronizing the onset of functional decline. As concentrations fall, only a subset of cells is positioned to regain activity, whereas under steady exposure each cell continues to decline over a broader temporal range according to its individual sensitivity. This history-dependent behavior mirrors the finding that transient responses in fluctuating environments determine long-term performance, a hallmark of fluctuation-adapted physiologies (13). Although our experiments were conducted *ex hospite* and therefore excluded host buffering and feedback processes, the findings suggest that environmental variability and cell-level heterogeneity interact to set resilience thresholds. Importantly, a stress spike is not inherently more damaging than constant exposure. Depending on how stress is structured in time, it may either amplify inhibition or create temporal windows that permit functional recovery, whereas constant stress suppresses recovery entirely. This adds a temporal dimension to classical dose–response theory and is consistent with theory and experiment showing that heterogeneous populations can achieve robustness in fluctuating environments (13, 23).

#### Outlook

Diuron provided a tractable PSII inhibitor for isolating temporal effects, but it represents only one of many chemical and environmental stressors experienced by *Symbiodiniaceae*. A key next step will be to explicitly test recovery following steady exposure, including how cells respond when stress is relaxed and whether prior exposure history conditions responses to subsequent steady or fluctuating perturbations. Extending similar analyses to temperature fluctuations, light variability, nutrient pulses, and multifactorial stress regimes will clarify whether the temporal sensitivities observed here reflect a broader physiological principle. Because environmental stress is inherently dynamic, the ability of microbial populations to persist may depend as much on *when* stress occurs as on its intensity. Applying comparable experimental frameworks to other systems, coupled with modeling of temporal response landscapes, may reveal general rules that link exposure structure, phenotypic heterogeneity, and resilience across microbial taxa and environments. By combining precisely timed perturbations with continuous single-cell measurements, this work outlines a mechanistic approach for predicting cellular outcomes under dynamic conditions and for understanding how exposure timing contributes to microbial resilience.

## Acknowledgments

This study was supported by grants from the Independent Research Fund Denmark (DFF-5281-00109B), the Swedish Research Council (2024-04748), the Novo Nordisk Foundation (NNF22OC0079370), the Science for Life Laboratory, and the Danish National Research Foundation (DNRF137) through the Center for Microbial Secondary Metabolites. We thank Luca Torello Pianale for insightful discussions and thoughtful feedback on data interpretation.

## Materials and Methods

### Materials

Polydimethylsiloxane (PDMS) elastomer (SYLGARD™ 184) was obtained from Dow Chemical. Ethanol (≥99.5%) was purchased from VWR International. Diuron (3-(3,4-dichlorophenyl)-1,1-dimethylurea; ≥98%) was obtained from Merck Life Science. A diuron stock (100 mg L⁻¹ in ethanol) was diluted into f/2 medium to prepare working solutions; carrier controls contained equivalent ethanol volumes.

### Microfluidic device fabrication

Microfluidic devices were fabricated in PDMS using standard photolithography (17). Microfluidic device designs have been previously published in (15) PDMS prepolymer (10:1 monomer:curing agent) was degassed, poured onto SU-8 master molds, and cured at 80 °C for 24 h. Devices were cut, inlet/outlet ports were punched (1.5 mm), and PDMS was plasma-bonded (Zepto 1, Diener electronics GmbH, Ebhausen, Germany) to cover slips (170 µm thickness) before baking them overnight at 80 °C. Devices were stored dust-free until use. Each microfluidic chip measured 15 × 7.2 mm and contained three parallel channels, each with a total of 4,200 microwells (20 µm diameter).

### Cell culture

*Fugacium* sp. (Clade F; CCMP2455) was obtained from the National Center for Marine Algae and Microbiota (Bigelow Laboratory). The strain originates from *Meandrina meandrites* (Jamaica) and was previously phylogenetically verified (14). Cells were maintained in f/2 medium (salinity 36 PSU, pH 8.0) prepared using artificial seawater (Instant Ocean). Medium was supplemented with nitrate (NaNO₃, 75 g L⁻¹), phosphate (NaH₂PO₄·H₂O, 5 g L⁻¹), silicate (Na₂SiO₃·9H₂O, 30 g L⁻¹), trace metals, and vitamins following the standard f/2 recipe. No antibiotics were used. Cultures were grown in 25 cm² culture flasks (Thermo Fisher Scientific) at 22°C under a 14:10 h light:dark cycle. White LED illumination (400–700 nm) was provided at 100 µmol photons m⁻² s⁻¹ using an AlgaeTron AG230 system (PSI, Czech Republic), and irradiance was verified with a spherical quantum sensor (US-SQS/L, Walz). All cultures were maintained in exponential phase and subcultured regularly to ensure physiological stability before experiments.

All work with *Fugacium* sp. was conducted under institutional biosafety level 1 (BSL-1) conditions in accordance with Uppsala University biosafety guidelines. The species is non-pathogenic and not subject to additional regulatory restrictions.

### Cell loading into microfluidic devices

Exponentially growing cultures (10–14 d post-inoculation) were centrifuged (1500 rpm, 2 min), resuspended in f/2 medium, and introduced into plasma-treated microfluidic devices. Microwell loading was enhanced by gentle compression of channel walls and gravitational settling for 1–2 h in the light incubator. Previous work demonstrated that this loading procedure does not affect Symbiodinaceae photophysiology (17). Excess cells were removed by flushing with fresh f/2 medium. Only single-cell wells were analyzed. Occupancy and cell position were verified using ImagingWin software.

### Chemical exposure of Symbiodiniaceae

Diuron exposures were conducted using a multichannel pressure pump (OB1 MK3+, ElveFlow, France) operated at a total flow rate of 60 µL min⁻¹. Flow stability, pulse fidelity and bubble removal were ensured using short microfluidic flow resistors (PEEK capillary tubing, 7.5 cm length, 100 µm ID), an inline microfluidic flow sensor (MFS3, 0-80 µL min⁻¹), and a PTFE membrane bubble trap (10 µm pore size, ElveFlow, France), respectively. All solutions were delivered in f/2 medium.

### Steady exposure profiles

For steady exposures, f/2 medium containing 4.3, 17.2, or 34.3 nM diuron was supplied continuously through a single pressurized inlet (700 mbar). These concentrations were selected based on preliminary assays demonstrating consistent, concentration-dependent reductions in *F_v_*/*F_m_* across the 24 h exposure window. Medium was prepared freshly for each experiment to minimize diuron photodegradation or adsorption losses.

### Fluctuating exposure profiles

Fluctuating diuron exposures followed a 24 h negative-cosine waveform, with peak concentrations of 9.2, 35.8, or 71.5 nM. The fluctuating profile was generated by driving two inlet channels, one containing diuron-free f/2 medium and one containing diuron-supplemented f/2, with opposing cosine pressure signals. The two streams were combined immediately upstream of the device via a Y-connector, producing a smoothly varying concentration profile inside the microfluidic channels (Figure 1a, S1).

To verify the dynamics of the imposed waveform, fluorescein was substituted for diuron and imaged under time-lapse fluorescence microscopy. The resulting intensity oscillations closely matched the programmed waveform (R² = 0.996), confirming that the delivered concentration profile accurately reproduced the intended fluctuations (Figure 1c, S2).

### Cumulative dose matching

To isolate the effects of timing from total exposure, steady and fluctuating treatments were designed to deliver identical cumulative chemical dose (nM·h) over both the first 12 h and the full 24 h exposure period. Cumulative dose was calculated as the time integral of the concentration profile,

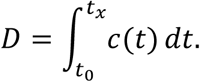

where *c(t)* is the instantaneous concentration and *t₀* and *tₓ* mark the start and end of the exposure window (12 h or 24 h). Time-averaged concentrations (4.3, 17.2, 34.3 nM) were used to label treatments, and peak concentrations for fluctuating profiles (9.2, 35.8, 71.5 nM) indicate the full range of stress experienced by cells.

### PAM imaging

Maximum quantum yield of PSII (*F_v_*/*F_m_* = (*Fₘ* − *F₀*)/*Fₘ*) was measured using a PAM imaging microscope (IMAG-RGB, Walz, Germany) equipped with a 10× Zeiss Fluar 10×/0.5 NA objective. Prior to each experiment, cells were dark-adapted for 15 min at 22 °C. Saturating pulses were applied every 15 min over the 24 h exposure period, and fluorescence images were acquired using ImagingWin software. This setup provides continuous single-cell photophysiological trajectories.

### Data analysis

Data analysis. Single-cell *F*ᵥ/*F*ₘ was quantified using PAMalysis (16) and analyzed in R (v4.3.1) and OriginPro (v10.2.0.196). Percent inhibition was calculated relative to time-zero values from unexposed controls. Cells with *F*ᵥ/*F*ₘ < 0.05 were classified as PSII-inactive. Cumulative dose for steady exposures was computed analytically; fluctuating exposures were integrated numerically from time-series concentration data. Heatmaps were generated using the pheatmap package, and K-means clustering (Euclidean distance) was used to partition cells into response groups based on their *F*ᵥ/*F*ₘ trajectories. Half-life values (t₁/₂) were extracted from smoothed single-cell trajectories as the time at which *F*ᵥ/*F*ₘ declined to 50% of its initial value. Experiments were performed across at least two independent biological replicates.

### Statistical analysis

Statistical analysis. Comparisons of single-cell *F*ᵥ/*F*ₘ values and t₁/₂ distributions between steady and fluctuating exposures were conducted using two-tailed Mann–Whitney U tests in OriginPro (v10.2.0.196). Statistical significance was defined as *p* < 0.05 unless otherwise indicated. To assess experiment-to-experiment variability, we compared experiment-averaged *F*ᵥ/*F*ₘ trajectories within each concentration; replicate trajectories were nearly identical, and no experiment effect was statistically resolvable. Summary statistics are reported as mean ± standard deviation. Cumulative dose calculations and visualizations were implemented in R.

### Data, materials, and software availability

All raw PAM fluorescence trajectories will be deposited in Dryad upon publication. Analysis scripts (R code for clustering, cumulative-dose calculations, and t₁/₂ extraction) will be made available on GitHub and archived in Zenodo with a DOI. Microfluidic device designs used in this study can be made available upon reasonable request. No new strains or unique biological materials were generated in this work.

### Replicates, randomization, and blinding

Experiments were performed across at least two independent biological replicates, each conducted on separate days with independently prepared cultures and microfluidic devices. Each biological replicate included hundreds of individual single-cell trajectories (*n* values reported in figures and legends). Because microwell trapping is deterministic and spatially defined, randomization and blinding were not applicable to these experiments.

## References

1. J. Wiedenmann, et al., Reef-building corals farm and feed on their photosynthetic symbionts. Nature 620, 1018–1024 (2023).

2. M. Stat, D. Carter, O. Hoegh-Guldberg, The evolutionary history of Symbiodinium and scleractinian hosts-Symbiosis, diversity, and the effect of climate change. Perspect Plant Ecol Evol Syst 8, 23–43 (2006).

3. O. Hoegh-Guldberg, Climate change, coral bleaching and the future of the world’s coral reefs. Mar Freshw Res 8, 839–866 (1999).

4. M. S. Roth, The engine of the reef: Photobiology of the coral-algal symbiosis. Front Microbiol 5, (2014).

5. J. W. van Dam, A. P. Negri, J. F. Mueller, R. Altenburger, S. Uthicke, Additive pressures of elevated sea surface temperatures and herbicides on symbiont-bearing foraminifera. PLoS One 7 (2012).

6. S. Morin, B. Chaumet, N. Mazzella, A time-dose response model to assess Diuron-induced photosynthesis inhibition in freshwater biofilms. Front Environ Sci 6 (2018).

7. M. Marzonie, et al., Toxicity thresholds of nine herbicides to coral symbionts (Symbiodiniaceae). Sci Rep 11 (2021).

8. S. E. Lewis, et al., Herbicides: A new threat to the Great Barrier Reef. Environmental Pollution 157, 2470–2484 (2009).

9. F. Flores, et al., Combined effects of climate change and the herbicide diuron on the coral Acropora millepora. Mar Pollut Bull 169 (2021).

10. P. Mercurio, et al., Degradation of herbicides in the tropical marine environment: Influence of light and sediment. PLoS One 11 (2016).

11. C. James, et al., Compilation of riverine water quality data from the Great Barrier Reef catchment area, northeastern Australia. Scientific Data 12 (2025).

12. J. Nguyen, J. Lara-Gutiérrez, R. Stocker, Environmental fluctuations and their effects on microbial communities, populations and individuals. FEMS Microbiol Rev 45 (2021).

13. J. Nguyen, et al., A distinct growth physiology enhances bacterial growth under rapid nutrient fluctuations. Nat Commun 12 (2021).

14. L. Xiao, et al., Photophysiological response of Symbiodiniaceae single cells to temperature stress. ISME J 16, 2060–2064 (2022).

15. M. Andersson, et al., A microscopy-compatible temperature regulation system for single-cell phenotype analysis – demonstrated by thermoresponse mapping of microalgae. Lab Chip 21, 1694–1705 (2021).

16. O. Pontén, et al., PACMan: A software package for automated single-cell chlorophyll fluorometry. Cytometry Part A 3, 203–213 (2023).

17. L. Behrendt, et al., PhenoChip: A single-cell phenomic platform for high-throughput photophysiological analyses of microalgae. Sci Adv 6, eabb2754 (2020).

18. A. Tlili, et al., Responses of chronically contaminated biofilms to short pulses of diuron. An experimental study simulating flooding events in a small river. Aquatic Toxicology 87, 252–263 (2008).

19. A. Negri, et al., Effects of the herbicide diuron on the early life history stages of coral in Marine Pollution Bulletin, (2005), pp. 370–383.

20. A. P. Negri, F. Flores, T. Röthig, S. Uthicke, Herbicides increase the vulnerability of corals to rising sea surface temperature. Limnol Oceanogr 56, 471–485 (2011).

21. J. W. van Dam, A. P. Negri, J. F. Mueller, S. Uthicke, Symbiont-specific responses in foraminifera to the herbicide diuron. Mar Pollut Bull 65, 373–383 (2012).

22. Joseph W. Bigger, TREATMENT OF STAPHYLOCOCCAL INFECTIONS WITH PENICILLIN BY INTERMITTENT STERILISATION. The Lancet 244, 497–500 (1944).

23. K. Lewis, Persister Cells. Annu Rev Microbiol 64, 357–372 (2010).

