## Supplemental materials for "Temporal structure of chemical stress controls single-cell inhibition and recovery in photosynthetic microorganisms"

Temporal stress structure, single-cell heterogeneity, microfluidics, Symbiodiniaceae, dynamic chemical exposure

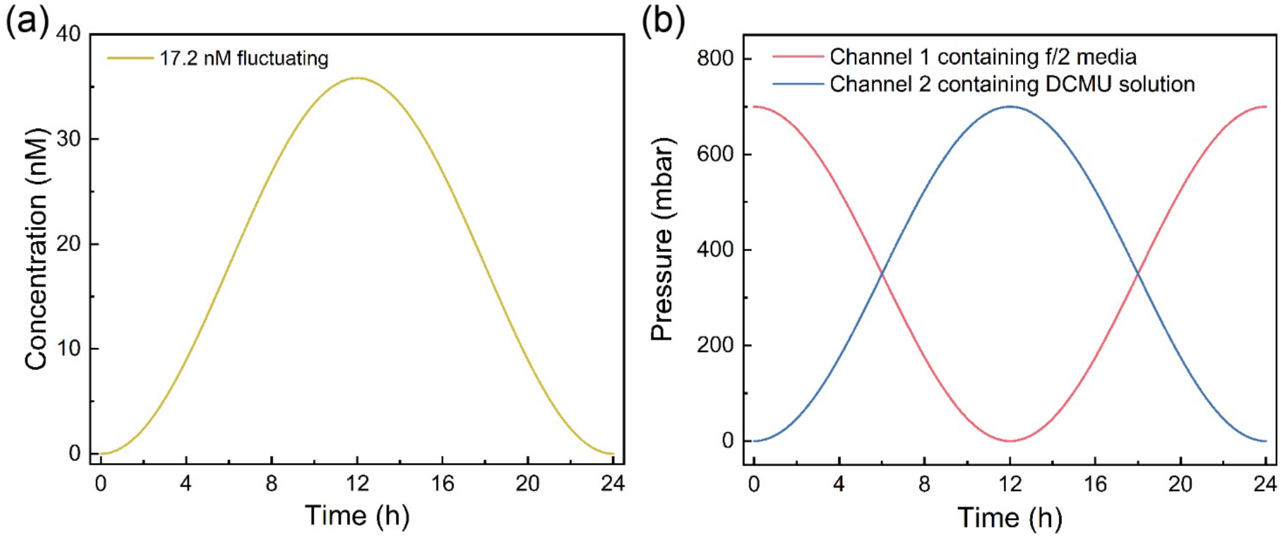


**Figure S1. (a)** Imposed fluctuating diuron concentration profile for the 17.2 nM time-averaged treatment. The waveform corresponds to a half-cycle of a negative cosine function with a peak concentration of 35.8 nM and a cumulative dose matched to the corresponding steady 17.2 nM exposure over 24 h. **(b)** Pressure profiles applied to inlet channels 1 and 2 to generate the fluctuating concentration profile. Channel 1 delivered a cosine-modulated pressure signal ranging from 700 to 0 mbar, while channel 2 delivered the inverse waveform (0 to 700 mbar). Their combination via a Y-junction produced the desired fluctuating diuron regime.

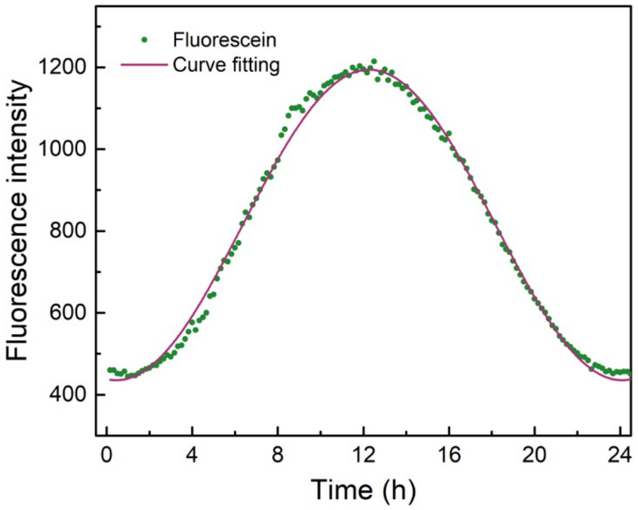


**Figure S2.** Fluorescein intensity measurements confirming the accuracy of the imposed fluctuating exposure profile. Green points show fluorescence intensity during the first half-cycle of the waveform, and the fitted sinusoidal curve (purple line) shows close agreement with the imposed signal ($R^2 = 0.996$). These results verify precise implementation of the fluctuating diuron regime inside the microfluidic device.

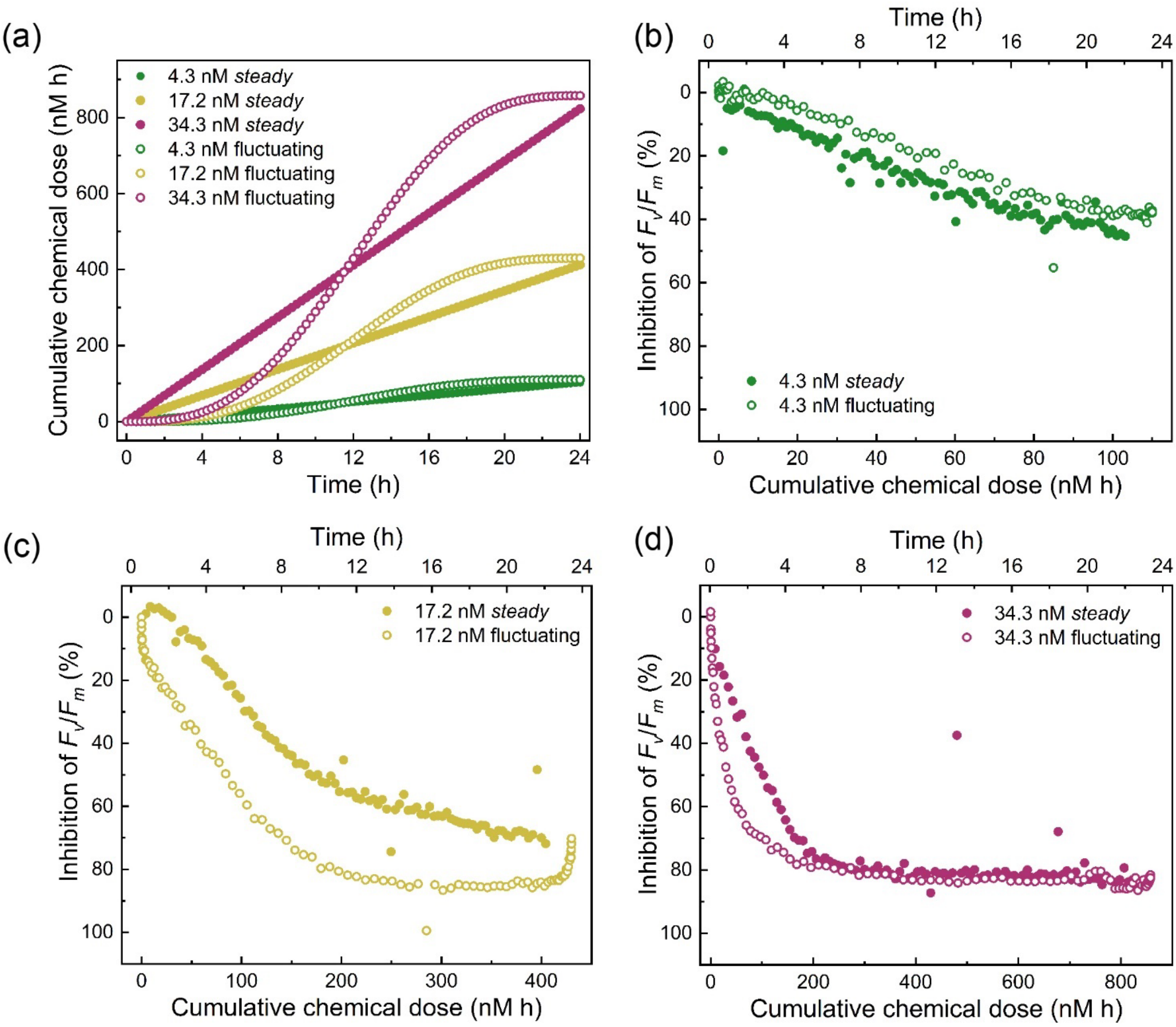


**Figure S3.** *(a)* Time-dependent cumulative diuron doses under steady and fluctuating exposure regimes at 4.3, 17.2, and 34.3 nM time-averaged concentrations. Fluctuating exposures delivered slightly higher cumulative doses (≈2–4%) than their steady counterparts due to minor mixing variability. *(b–d)* Inhibition of mean $F_v/F_m$ as a function of experimentally measured cumulative dose for steady (closed circles) and fluctuating (open circles) exposures at *(b)* 4.3 nM, *(c)* 17.2 nM, and *(d)* 34.3 nM. These plots show that differences in inhibition between regimes are consistent with effects of exposure timing rather than cumulative dose alone.

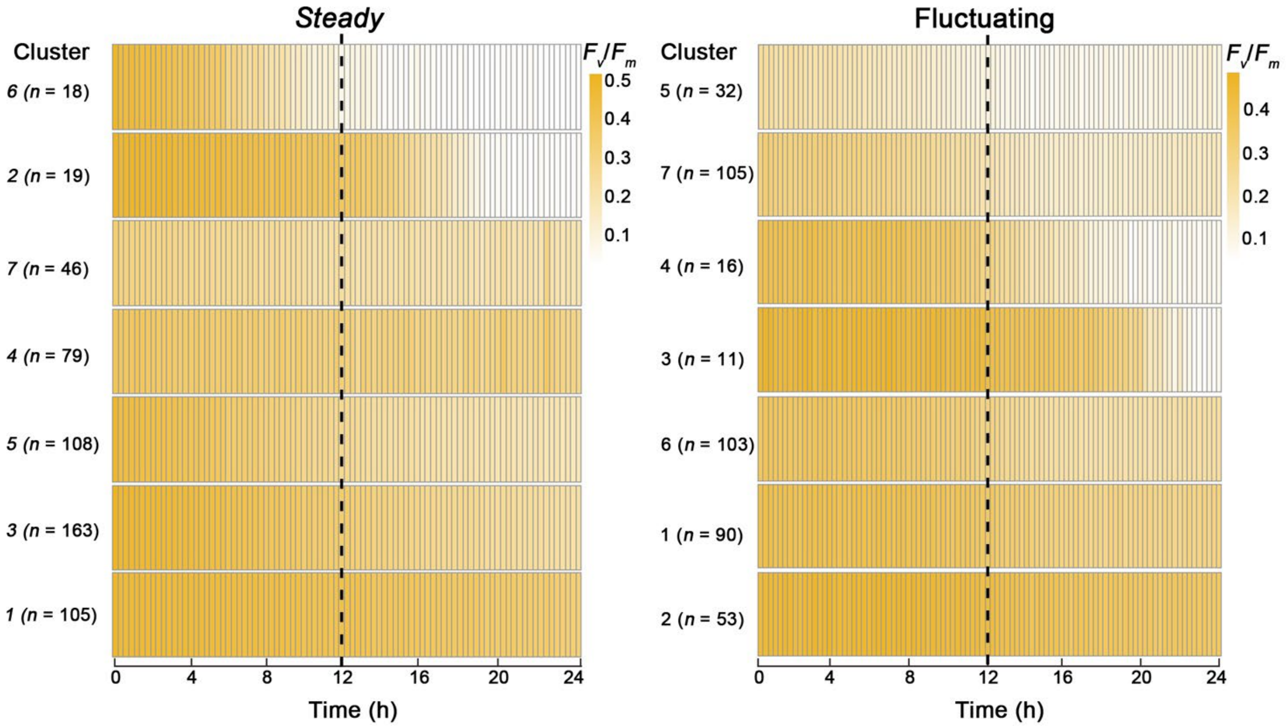


**Figure S4.** Single cell $F_v/F_m$ trajectories at 4.3 nM diuron resolved by K-means clustering. Heatmaps show individual cell trajectories under steady (left) and fluctuating (right) exposure regimes. Each horizontal block represents a distinct response cluster, with *n* indicating the number of cells assigned to that cluster. Columns represent time points over the 24-h experiment, and colors denote instantaneous $F_v/F_m$. The vertical dashed line marks the 12-h midpoint at which cumulative dose is matched between regimes. Even at low stress, cells occupy distinct functional trajectories that differ in the magnitude and timing of inhibition.

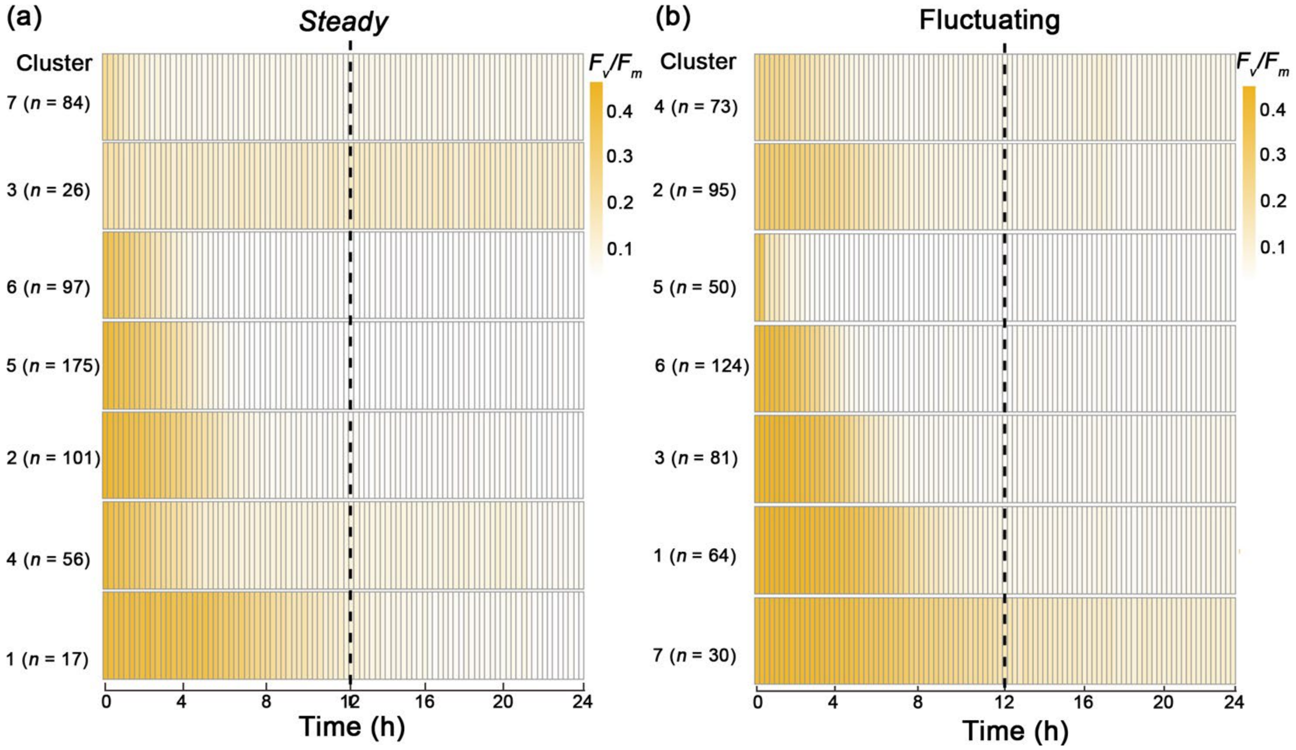


**Figure S5**. Single-cell $F_v/F_m$ trajectories at 34.3 nM diuron resolved by K-means clustering. Heatmaps show individual cell trajectories under steady *(a)* and fluctuating *(b)* exposures. Each block represents one response cluster, with *n* indicating the number of cells in that cluster. Colors represent instantaneous $F_v/F_m$ over the 24-h period. The dashed line marks the 12-h cumulative-dose match point. At this high stress level, clustering reveals strong functional differentiation, including cells that maintain residual activity, rapidly lose function, or (under fluctuating exposure) show transient late-stage increases in activity before complete inhibition.

**Table S1.** Inhibition of mean $F_v/F_m$ (%) after 12 h and 24 h under steady and fluctuating diuron exposures. Values represent population-averaged inhibition relative to unexposed controls for each time-averaged concentration (4.3, 17.2, and 34.3 nM). For fluctuating regimes, peak concentrations were 9.2, 35.8, and 71.5 nM, respectively

| Inhibition of average $F_v/F_m$ (%) | 12 h | 24 h |
|---|---|---|
| *Steady* 4.3 nM | 26.3 ± 18.3% | 45.3 ± 19.4% |
| Fluctuating 4.3 nM | 19.2 ± 3.6% | 37.7 ± 27.4% |
| *Steady* 17.2 nM | 55.7 ± 30.8% | 60.5 ± 3.1% |
| Fluctuating 17.2 nM | 82.3 ± 11.0% | 70.2 ± 11.8% |
| *Steady* 34.3 nM | 100% | 100% |
| Fluctuating 34.3 nM | 100% | 100% |
